# Spatial organization of the kelp microbiome at micron scales

**DOI:** 10.1101/2020.03.01.972083

**Authors:** S. Tabita Ramirez-Puebla, Brooke L. Weigel, Loretha Jack, Cathleen Schlundt, Catherine A. Pfister, Jessica L. Mark Welch

## Abstract

Macroalgae are colonized by complex and diverse microbial communities that are distinct from those on inert substrates, suggesting intimate symbioses that likely play key roles in both macroalgal and bacterial biology. Canopy-forming kelp fix teragrams of carbon per year in coastal kelp forest ecosystems, yet little is known about the structure and development of their associated microbial communities. We characterized the spatial organization of bacterial communities on blades of the canopy-forming kelp *Nereocystis luetkeana* using fluorescence *in situ* hybridization and spectral imaging with a probe set combining phylum, class and genus-level probes to target >90% of the microbial community. We show that kelp blades host a dense microbial biofilm, generally less than 20 μm thick, in which disparate microbial taxa live in close contact with one another. The biofilm is spatially differentiated, with tightly clustered cells of the dominant symbiont *Granulosicoccus sp*. (Gammaproteobacteria) close to the kelp surface and filamentous *Bacteroidetes* and Alphaproteobacteria relatively more abundant near the biofilm-seawater interface. Further, a community rich in *Bacteroidetes* colonized the interior of kelp tissues. Microbial community structure and cell density increased along the length of the kelp blade, from sparse microbial colonization of newly produced tissues at the meristematic base of the blade to an abundant microbial biofilm on older tissues at the blade tip. Finally, kelp from a declining population hosted fewer microbial cells compared to kelp from a stable population, indicating that biofilms are characteristic of health and that biofilm loss may be related to the condition of the host.

**Importance:** The microbial community coating the surfaces of macroalgae may play a key but underexplored role both in the biology of the macroalgal host and in the biogeochemistry of the coastal ocean. We show that photosynthetic blades of the canopy-forming kelp *Nereocystis luetkeana* host a complex microbial biofilm that is both dense and spatially differentiated. Microbes of different taxa are in intimate cell-to-cell contact with one another; microbial cells invade the interior of kelp cells as well as cover their external surfaces; and a subset of the surface microbiota projects into the water column. These results highlight the potential for metabolic interactions between key members of the kelp microbiome as well as between microbes and their host. The dense layer of microbes coating the surface of the kelp blade is well-positioned to mediate interactions between the host and surrounding organisms and to modulate the chemistry of the surrounding water column.

## Introduction

Macroalgae are foundational members of their local ecosystems, where they provide animal habitat and nursery areas (Lamy et al. 2020), contribute to primary productivity (Wilmers et al. 2012), and modify surrounding seawater chemistry (Pfister et al. 2019). The surfaces of macroalgae are associated with microbial communities that may play a key but underexplored role in macroalgal biology. The surface of macroalgae is frequently colonized by a microbial community that is spatially well-positioned to act as a mediator of algal metabolic exchange with the environment. As dominant members of temperate coastal oceans, brown algae known as kelp host microbial communities distinct from those in surrounding seawater (Michelou et al. 2013, Chen & Parfrey 2018, Weigel and Pfister 2019) and from rocky substrates (Lemay et al. 2018), indicating that kelp may possess mechanisms for selecting or recruiting a unique subset of water column microbes, while preventing fouling or biofilm establishment of many others. Further, kelp metagenomes contain a high abundance of microbial motility genes (Minich et al. 2018) and kelp tissue only several days old becomes colonized by bacteria (Weigel and Pfister 2019). Microbial community changes have been linked to algal host health (Marzinelli et al. 2015) and environmental stressors (Minich et al. 2018). While the functions of macroalgal microbiomes are still being elucidated, surface-associated microbes can metabolize algal polysaccharides (Martin et al. 2015, Lin et al. 2018) and in seagrasses, functionally important metabolite exchanges between host and microbes have been demonstrated (Tarquinio et al. 2018). Despite the potential for macroalgal-associated microbial communities to affect nutrient exchange, biofouling, disease and even host development (Egan et al. 2013), we know little about the composition and development of macroalgal microbial communities.

Imaging of microbial community organization reveals the micrometer-scale localization of taxa relative to one another and relative to host tissue and other landmarks such as the surface of the biofilm. In a complex microbial community characterized by cross-feeding and metabolic interactions among diverse partners, localization provides clues about the micro-habitats and metabolic partners that foster the growth of particular microbes. Localization is key because microbes interact primarily with other microbes within a distance of a few microns or tens of microns, particularly in environments characterized by fluid flow (Kolenbrander et al. 2010, Cordero and Datta 2016, Dal Co et al. 2019). Thus, visualizing the spatial structure of a microbial biofilm contributes greatly to our understanding of host-microbe and microbe-microbe interactions.

We investigated the micron-scale spatial organization of microbial communities living on photosynthetic blades of bull kelp, *Nereocystis luetkeana*. Bull kelp are a highly productive component of the northeast Pacific Ocean, creating vast underwater forests from southern California to Alaska. This annual kelp displays extraordinarily high growth rates, with photosynthetic kelp blades growing outwards from the kelp thallus at rates of 0.5 – 2 cm per day (Weigel and Pfister 2019) and reaching heights greater than 40 m. This rapid growth permitted us to ask how microbial community spatial structure and diversity changes from newly produced tissue at the base of the blade to months-old tissue at the blade tip. We used Combinatorial Labeling and Spectral Imaging – Fluorescence *in situ* Hybridization (CLASI-FISH; Valm et al. 2011, 2012) to investigate microbial community spatial structure on *N. luetkeana* blades during its rapid summer growth. In addition, we compared the microbial biofilm structure from a healthy *N. luetkeana* population to a geographically distinct population that has been in decline in recent years (Pfister et al. 2017, Berry et al. 2020). We demonstrate that the composition of the kelp microbiome displays repeatable spatial structure (i.e. microbiota changes over micron-scale distances from the blade surface), that colonization density correlates with the age of the blade, and that bacterial diversity and density are related to the state of health of the kelp.

## Results

### Development of probe set and sample preparation methodology for CLASI-FISH on kelp

To investigate the community structure of the kelp microbiota over time and in kelp populations at different locations in Washington State, we collected kelp blade tissue samples (n = 6 individuals) from two sites reported in Weigel and Pfister (2019): Squaxin Island, in Southern Puget Sound, and Tatoosh Island, on the outer coast of the Olympic Peninsula. In this study, we carried out CLASI-FISH and imaging on a total of 15 samples: 3 from a declining population at Squaxin Island (Berry et al. 2020) and 12 from a persistent population at Tatoosh Island (Table S1). Squaxin Island kelp were sampled once on 21 June 2017, while on Tatoosh Island we collected a time series consisting of 5 collection days from June to August 2017 (Table S1), spanning the peak in annual biomass. The 12 imaged samples from Tatoosh Island include 4 pairs (n = 8 samples) from both the base (near the meristematic tissue) and the tip (older tissue) of the same kelp frond over the time series, and an additional 4 samples from the tip of different kelp blades in June and July (Table S1). Samples were processed for 16S rRNA gene sequencing as previously reported (Weigel and Pfister 2019); additional samples, taken in parallel, were collected and preserved in 95% ethanol for CLASI-FISH.

Characterizing the microbiome of *N. luetkeana* with 16S rRNA gene sequencing revealed a community that was diverse but composed of the same major taxa at all sampled time points and both locations (Fig. 1, Weigel and Pfister, 2019). The most abundant genus-level taxon was *Granulosicoccus* (Gammaproteobacteria), which accounted for close to half of the sequences overall in samples from Tatoosh Island (Fig. 1); other major taxa included Alphaproteobacteria, Bacteroidetes, and Verrucomicrobia. In samples from the declining Squaxin Island kelp population, Alphaproteobacteria were dominant, comprising > 75% of the microbial community (Fig. 1).

**Figure 1.**
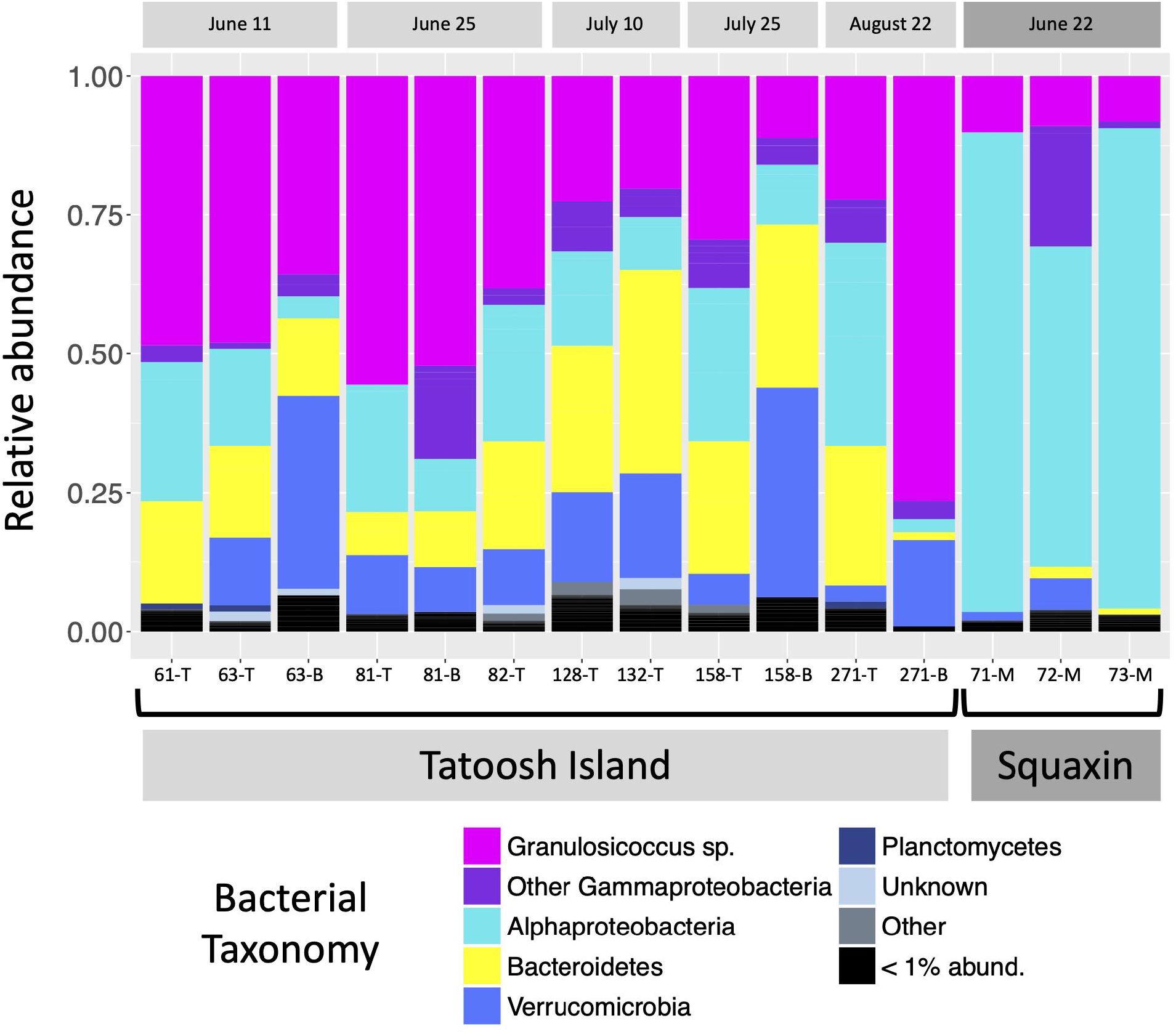
Relative abundance of bacterial taxa on the kelp *N. luetkeana*, grouped to show taxa detected by our CLASI-FISH probe set. 16S rRNA gene sequencing showed that bacterial composition is broadly consistent throughout the summer. The most abundant taxa were Verrucomicrobia, Bacteroidetes, Alphaproteobacteria and Gammaproteobacteria. The gammaproteobacterial genus *Granulosicoccus* was in high abundance in samples from Tatoosh Island. In samples from a declining population at Squaxin Island *Granulosicoccus* spp. were still present but Alphaproteobacteria were dominant. Collection date is shown at top. Collection site is shown at bottom. Sample number and part of the kelp blade sampled are indicated below each column. T= tip; B= base; M= middle.

In contrast to sequencing, which provides information on community structure at centimeter scales, imaging permits micrometer-scale analysis of microbial spatial organization. To investigate the spatial organization of the kelp microbiota we developed a probe set for CLASI-FISH (Combinatorial Labeling and Spectral Imaging- Fluorescence *in situ* Hybridization) to enable simultaneous identification and imaging of the major bacterial groups. For comprehensive coverage of bacteria, we used the probes Eub338-I, II, and III (Amann et al. 1990, Daims et al. 1999) using one fluorophore for probe Eub338-I and a different fluorophore to label both Eub338-II and Eub338-III, to differentiate the Verrucomicrobia and Planctomycetes from the rest of the Bacteria. For greater taxonomic resolution within the bacteria identified by Eub338-I, the probe set contained probes specific for the major groups comprising the *N. luetkeana* microbiota: Bacteroidetes, Alphaproteobacteria, and Gammaproteobacteria. For added taxonomic resolution within Gammaproteobacteria we designed two new probes targeting the most abundant genus, *Granulosicoccus*. Thus, the probe set targeted nested levels of taxonomic identification, with cells identified by combinations of one, two, or three fluorophores (Table 1). We tested probe specificity by applying the set of 7 probes to 5 pure cultures. Each probe hybridized with its target taxa and only faint hybridization was observed with nontarget taxa (Fig. S1 and S2).

**Table 1.**
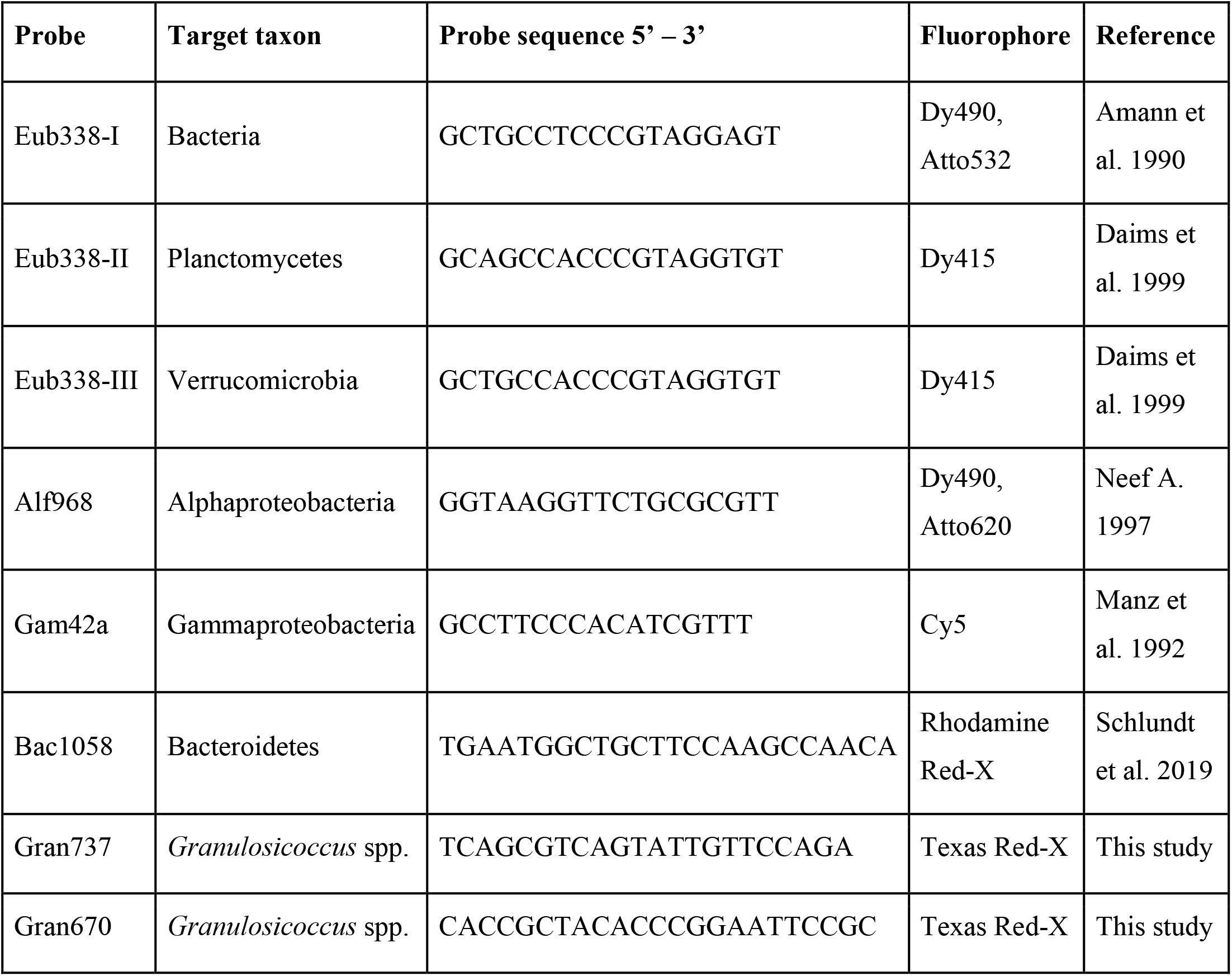
Probes used in this study.

For sample preparation we employed two strategies: whole-mount preparation and embedding and sectioning. Whole-mount preparations enable imaging of a large surface area of kelp blade but have the disadvantage that layers of the kelp tissue tend to separate over the course of the hybridization and washing steps. We found that coating the tissue with a layer of agarose at the beginning of the experiment helped the sample to remain intact during subsequent manipulations (see Methods). An alternative preparation procedure, in which the sample was embedded in methacrylate resin followed by sectioning and application of probes to the sections mounted on a slide, preserved spatial organization by immobilizing the sample in resin and permitted imaging of thin cross-sections through the kelp blade.

Imaging of such cross-sections of kelp embedded in methacrylate enabled us to visualize the overall relationship of the microbiota to the underlying kelp tissue (Fig. 2A). The kelp tissue itself is visible in a transmitted-light image as a series of large irregularly shaped chambers (sieve tubes), with a row of oblong photosynthetic cells along both the upper and the lower surface of the blade (Fig. 2A). The microbes are located primarily in a dense layer several micrometers thick on the exterior of both the upper and lower surface. Imaging at a magnification sufficient to visualize individual bacterial cells shows each taxon as single cells or small clumps, often immediately adjacent to cells of different taxonomic identification (Fig. 2A, insets i-iii). In addition to the surface layer, some bacteria are present in the interior of the blade; a subset of the community consisting largely of Bacteroidetes rods can be seen within the kelp tissue immediately beneath the surface layer (Fig. 2Ai), and a small cluster of mixed composition is visible near the center of the image (Fig. 2Aii). Thus, a variety of modes of interaction between microbiota and host can be visualized in cross-sectional view.

**Figure 2.**
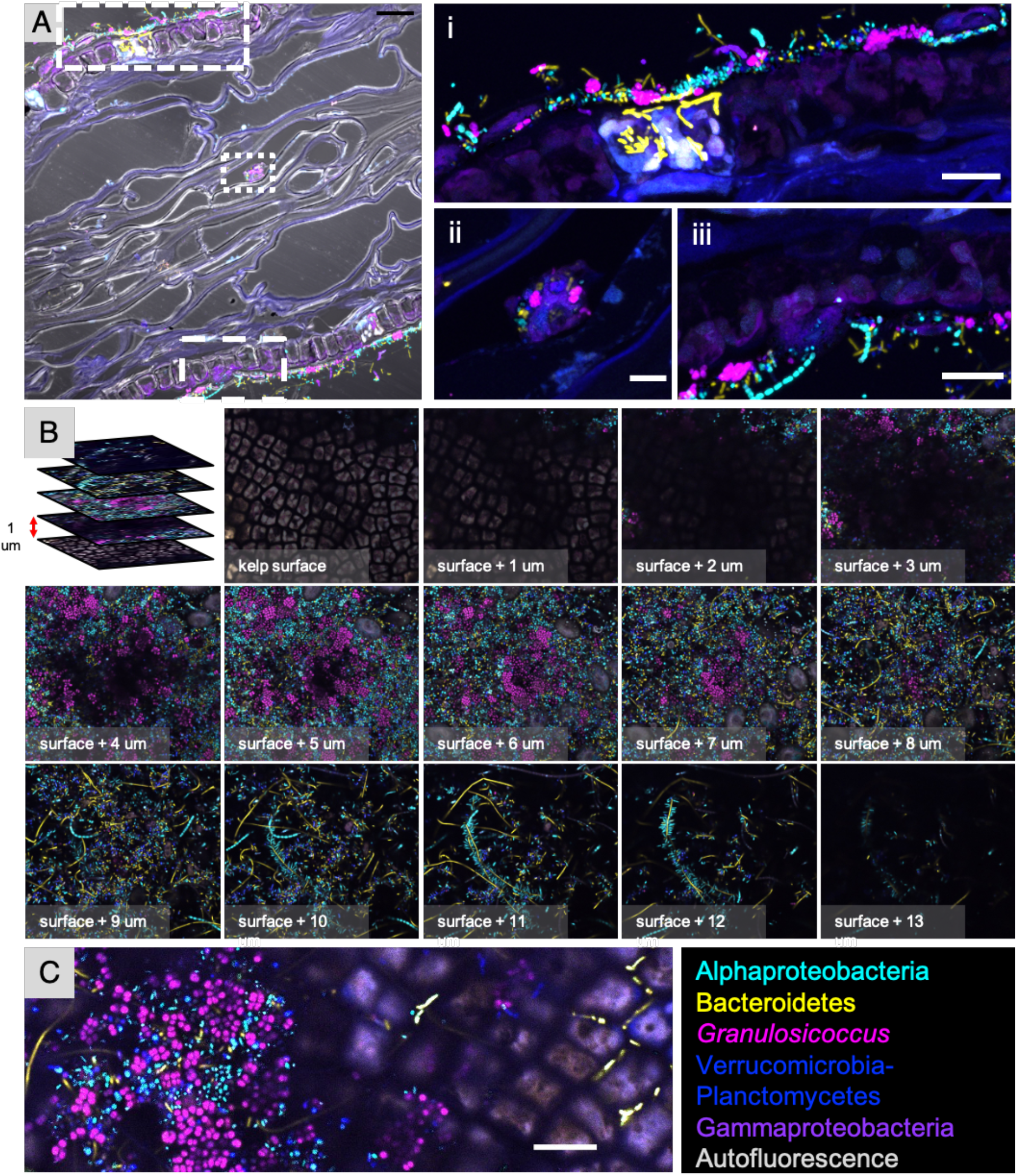
Cross-section, whole-mount, and oblique optical section images give different views of the biofilm on blades of *N. luetkeana*. Kelp blades were subjected to FISH with a probe set for the 5 major bacterial groups. (A) Cross-section image of a kelp blade embedded in methacrylate. Merge of transmitted light and confocal images shows biofilm on both surfaces and some colonization in center. (i), (ii) and (iii) are enlarged images of the dashed rectangles in panel (A). Bacteroidetes rods present within the autofluorescent region of kelp tissue in panel (i) fluoresce more brightly than rods in the surface biofilm and therefore appear overexposed in the image. (B) Whole-mount preparation imaged as a z-stack; planes 1 micrometer apart in the z dimension are shown. (C) Oblique optical section showing biofilm at left and kelp surface at right. Bacteroidetes rods are visible between kelp surface cells (right). Scale bar = 20 μm in (A); 5 μm (i), (ii) and (iii); 10 μm in (C).

By contrast, placing a whole-mount of a kelp blade flat on a microscope slide permitted imaging of a vertical series (or z-stack) of images from the surface of the kelp blade through the biofilm (Fig. 2B). This whole-mount imaging reveals the relationship of microbial cells to one another and the changing composition of the community as a function of distance from the kelp blade. Moving up from the plane in which the kelp photosynthetic cells are visible (Fig. 2B), serial optical sections show first a largely fluorescence-free region a few micrometers thick and then dense colonization by a mixed microbial community. At approximately 3 to 6 μm from the surface of the blade, clusters of *Granulosicoccus* cells are prominent and diatoms are visible (Fig. 2B). In the region approximately 8 to 12 μm from the blade surface, filaments are a more prominent part of the community. Non-filamentous cells of Alphaproteobacteria, Bacteroidetes and Verrucomicrobia or Planctomycetes are present throughout the biofilm. The whole-mount images, like the cross-sections, show that cells of diverse taxa are directly adjacent to one another in the kelp surface biofilm.

In perfectly flat whole-mount images one sees either the algal surface or the microbiota but not both unless the microbiota has invaded into the tissue. In many whole-mount images, however, the algal surface and the overlying microbiota can be seen simultaneously because the sample is tilted relative to the plane of imaging, such that the confocal microscope image is an optical section through the sample at an oblique angle and a single plane of focus captures both the kelp blade and the microbial community. In the example in Fig. 2C, kelp cells are visible as large (∼5 μm × 10 μm) oblongs at the right; Bacteroidetes spp. are intercalated between the kelp cells. In the center of the image a scattering of taxonomically mixed bacteria are located on or between the kelp cells, while at the left-hand side of the image the full microbial community is visible. As most whole-mount preparations are not entirely flat, many images represent to one degree or another an oblique-angle view of the material.

### Bacteria on kelp blades form mixed epiphytic communities whose abundance is dependent on the age of the underlying tissue

Due to the high growth rate of the kelp thallus, producing multiple centimeters of new tissue in a single day (Weigel and Pfister 2019), it is possible to study how spatial structure and diversity of the microbial community differ between newly produced tissue vs. tissue that is weeks to months old. Although the microbial community on older tissue contains more microdiversity, the composition and relative abundances of major taxa are broadly consistent across young and old tissues (Fig. 1, Weigel and Pfister 2019). However, imaging showed a higher density of colonization at the tip of the blade, which is months old, compared to the base, where newly produced tissue is only days old (Fig. 3). At the tip (Fig. 3 A-C) cells form a confluent biofilm, whereas at the base only scattered cells are visible (Fig. 3 D-F). This pattern is observed in samples collected throughout the summer (Fig 3).

**Figure 3.**
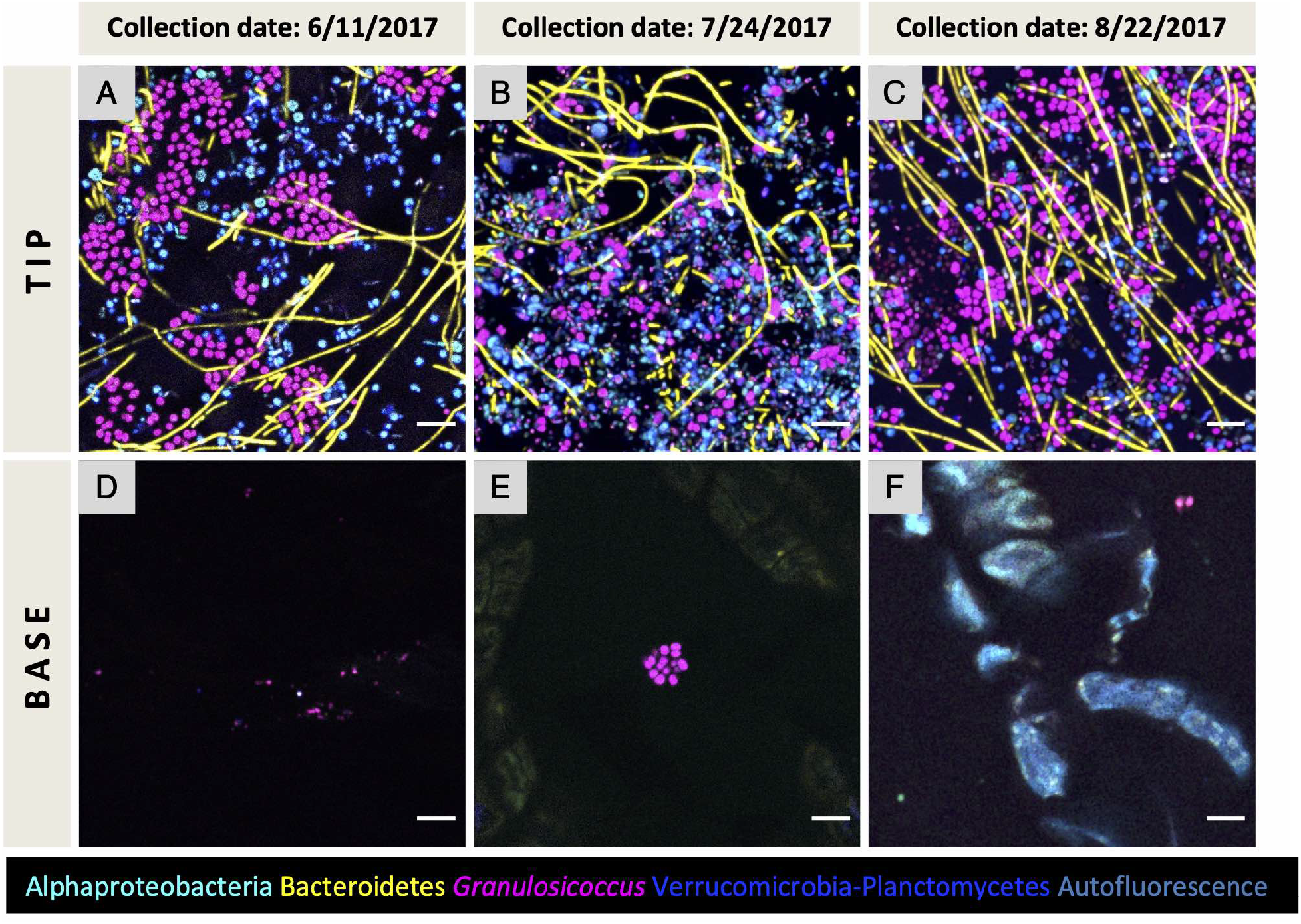
Bacterial abundance of the surface biofilm in old and young tissue. Whole-mount images of tip and base tissues collected from the same kelp frond. Three individuals from different collection dates are shown. Older tissue of the tip (A, B, C) is densely colonized compared to young tissue of the base (D, E, F). The same pattern is observed throughout the summer. Scale bar= 5 μm (A-F).

Typical images of the tip community (Fig. 4) show that bacteria at the tip of blades are mixed at micron scales. *Granulosicoccus*, the most abundant genus in the 16S rRNA gene sequencing, formed patches or clusters up to 15 μm in diameter in some samples (Fig. 4A); close inspection reveals cells of other taxa nestled within the clusters (Fig. 4A). Alphaproteobacteria, Bacteroidetes rods, Verrucomicrobia and Planctomycetes did not form large clusters but instead are intermixed in the biofilm. In whole-mount preparations, Bacteroidetes filaments lie on top of the other bacteria (Fig. 4A, B), suggesting that they are located in a different level of the biofilm and become flattened onto the sample during the FISH and mounting procedure. This observation is reinforced by cross-section images, which show a biofilm typically 3 to 5 μm thick, with filaments projecting approximately 10 μm from the biofilm (Fig. 4C).

**Figure 4.**
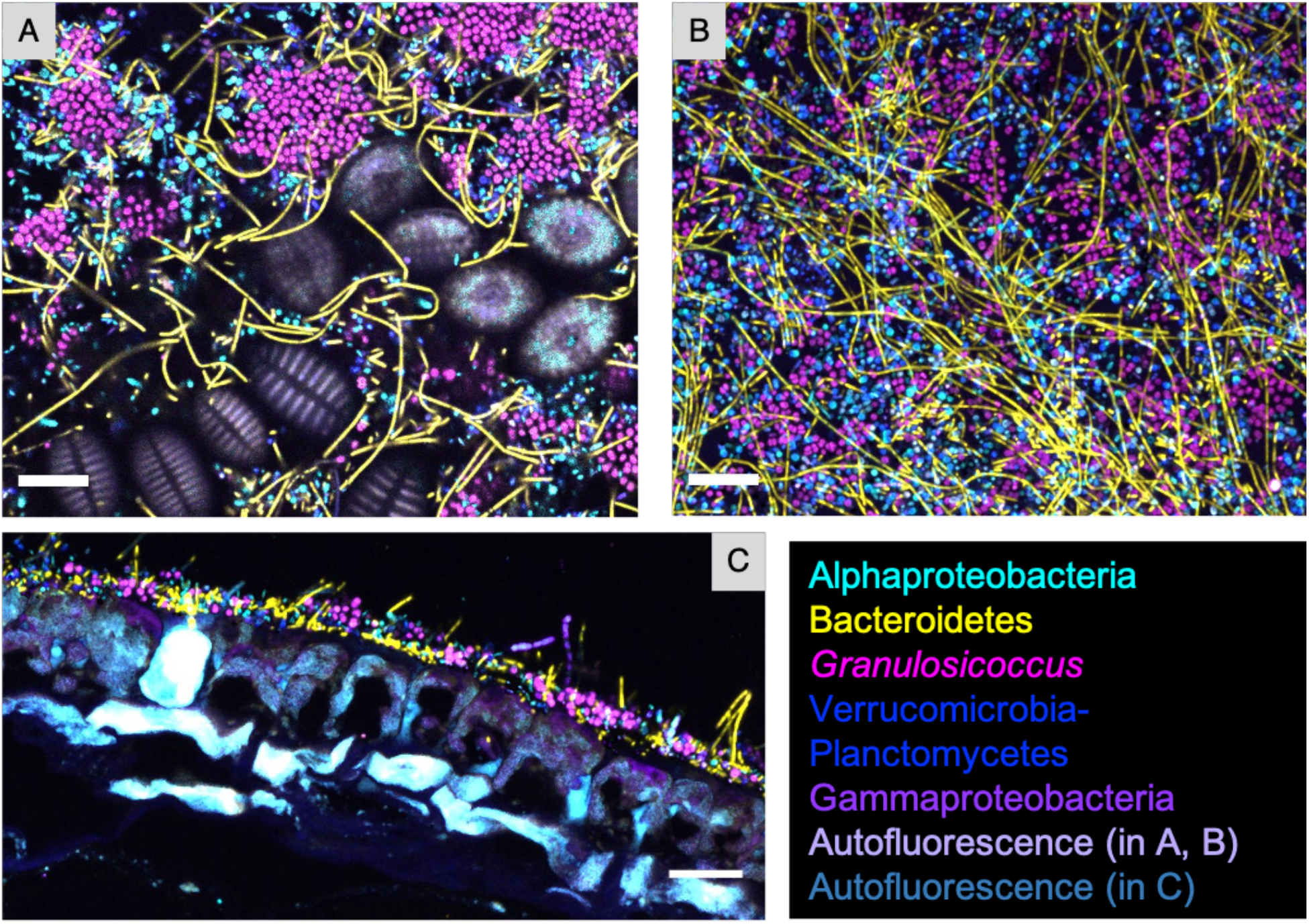
Spatial structure of the epiphytic microbial community at the tip of *N. luetkeana* blades. Bacteria at the tip of kelp blades form a dense biofilm. (A) and (B) show whole-mount images of samples collected in different dates; (C) is a cross-section showing thickness of the surface biofilm. Microorganisms are intermixed, often directly adjacent to cells of disparate taxa and always within 10 microns of other taxa. *Granulosicoccus* aggregate in clusters while other taxa are more dispersed. Abundant Bacteroidetes filaments appear to be lying on the other taxa in the whole mount and are visible in the cross-section, together with filaments of Gammaproteobacteria, projecting into the water column. Diatoms surrounded by bacteria are visible in (A). Scale bar= 10 μm (A), (B) and (C).

In addition to a dense bacterial biofilm, colonies of diatoms were observed, mainly in samples collected in June and July 2017 (Fig.4A; Fig. 5). In a cross-sectional image, it was apparent that diatoms were embedded within the biofilm and a layer of bacteria is observed between the diatoms and the kelp tissue (Fig. 5C, D), suggesting that the diatoms colonized the host after the bacteria. No specific association between diatoms and particular bacterial taxa was observed.

**Figure 5.**
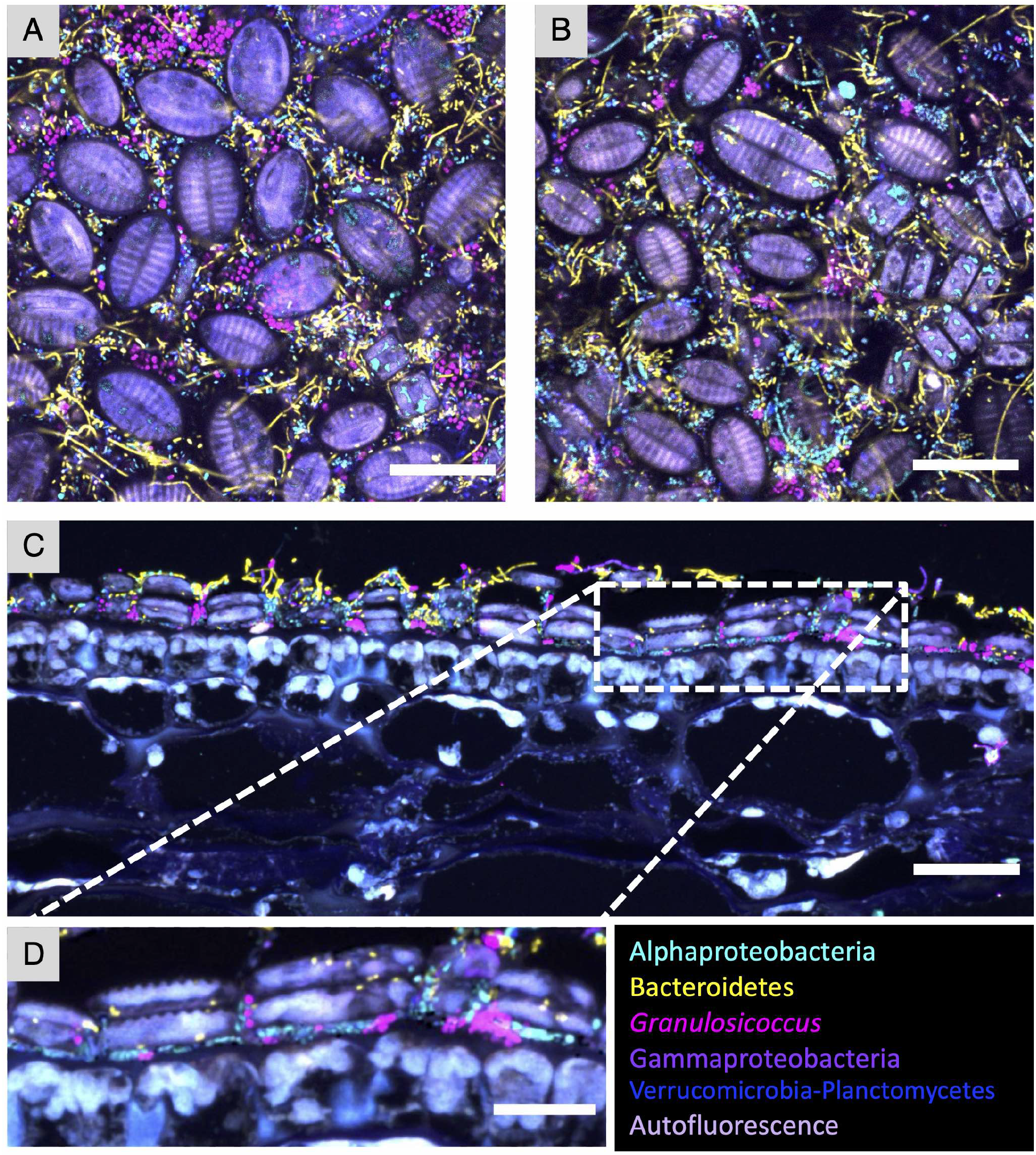
Diatoms are part of the microbial community of *N. luetkeana*. Colonies of diatoms were observed on kelp blades in samples from June and July. (A) and (B) Representative images of whole-mount FISH showing diatoms surrounded by bacteria. (C) Cross-section showing diatoms embedded within the bacterial biofilm. (D) Enlarged image of the dashed rectangle in (C). No specific association between diatoms and particular taxa was observed. Scale bar= 20 μm (A-C); 10 μm (D).

### Endophytic bacteria and direct interactions with the blade surface

Endophytic bacteria have been reported in macroalgae (Hollants et al. 2011) but have not previously been imaged with probes that could distinguish multiple taxa. In addition to the superficial biofilm, we detected bacteria inside kelp tissue by imaging cross-sections (Fig. 6). Bacteroidetes rods and, less abundantly, other members of the community were observed colonizing intercellular spaces of the outermost layer of kelp cells in samples from July (Fig. 2A,2C, Fig. 6). These endophytic Bacteroidetes were located adjacent to kelp cells that may be especially metabolically active given their strong autofluorescence. Microbes occasionally colonized the surface layer of cells directly (Fig. 6B), but in contrast to the Bacteroidetes-rich invasion into highly autofluorescent regions, no obvious preference for any underlying morphology of the kelp was detected. We also observed the microbial community forming small clusters on the interior of the frond (Fig. 2Aii) or forming a strand running through the kelp inner tissue (Fig. 6C, D).

**Figure 6.**
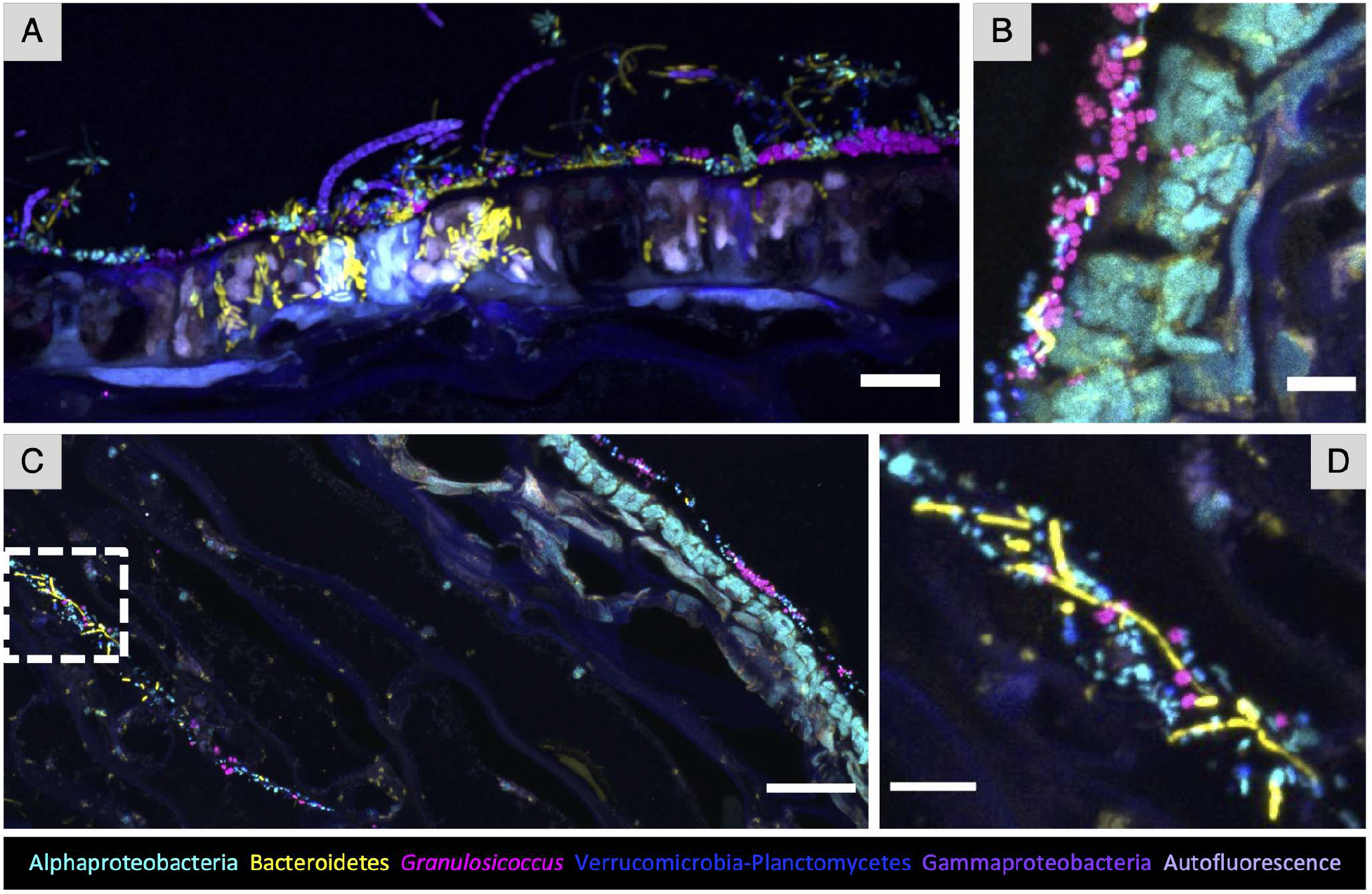
Endophytic bacteria of *N. luetkeana*. (A) Cross-section showing Bacteroidetes rods colonizing intercellular spaces of brightly autofluorescent kelp surface cells. (B) A region in which the biofilm is directly adjacent to the kelp tissue and some *Granulosicoccus* are observed between kelp cells. (C) Bacteria were also detected colonizing deeper areas of the tissue, in this instance around 120 μm from the surface. (D) enlarged image of the dashed rectangle in (C). Scale bar= 10 μm (A); 5 μm (B) and (D); 20 μm (C).

### Unipolar labeling of adherent Alphaproteobacteria with wheat germ agglutinin

Wheat germ agglutinin (WGA) is a lectin that binds to N-acetylglucosamine and N-acetylmuramic acid residues that can be present both in bacterial cell walls and in host mucus secretions. We stained samples with fluorophore-labeled WGA and observed staining in spots on the kelp tissue itself, in cells hybridizing with the *Granulosicoccus* probe, and most intriguingly, on cells hybridizing with the *Alphaproteobacteria* probe that reacted asymmetrically with the WGA, showing fluorescence at only one end of the cell (Fig. 7). This pattern was observed across multiple kelp collected in different months and sites (Fig. 7A, B) including the declining population of kelp from Squaxin Island (Fig. 8D, E). In cross-section images the exopolysaccharides were present at the end of the cell that was in contact with the kelp surface (Fig. 7C) suggesting a potential role of the WGA-stained structure in adhesion of the microbe to the kelp.

**Figure 7.**
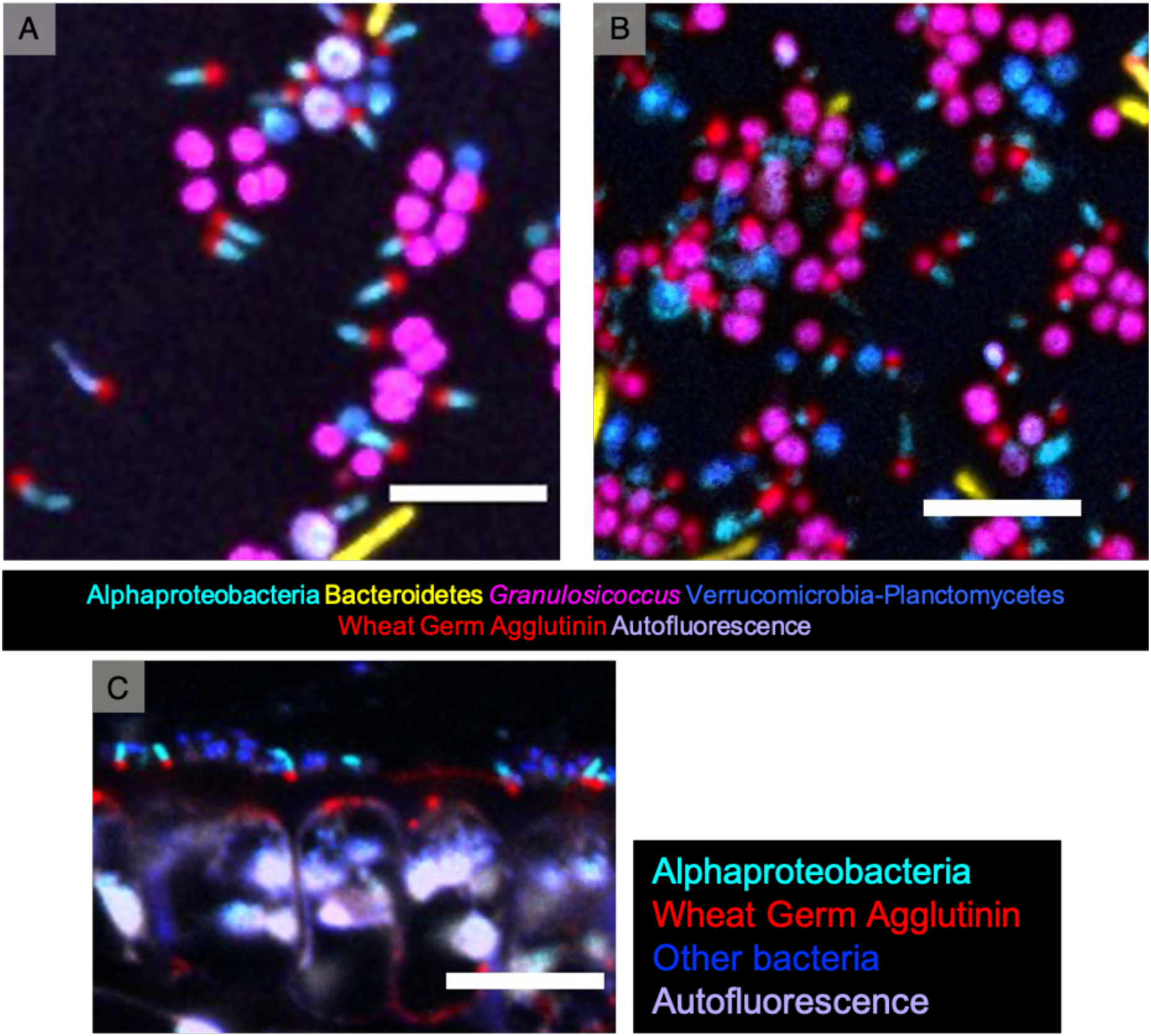
Unipolar labeling of adherent Alphaproteobacteria by wheat germ agglutinin. Wheat germ agglutinin was used to stain N-acetylglucosamine and N-acetylmuramic acid residues. Staining was observed on Alphaproteobacteria rods at only one end showing apparent polarity with respect to the cells. (A) and (B) Representative images of whole-mount FISH in samples collected in different months. (C) Cross-section image showing Alphaproteobacteria rods with the polar polysaccharide end attached to the kelp surface. Scale bar=5 μm (A) and (B); 10 μm (C).

**Figure 8.**
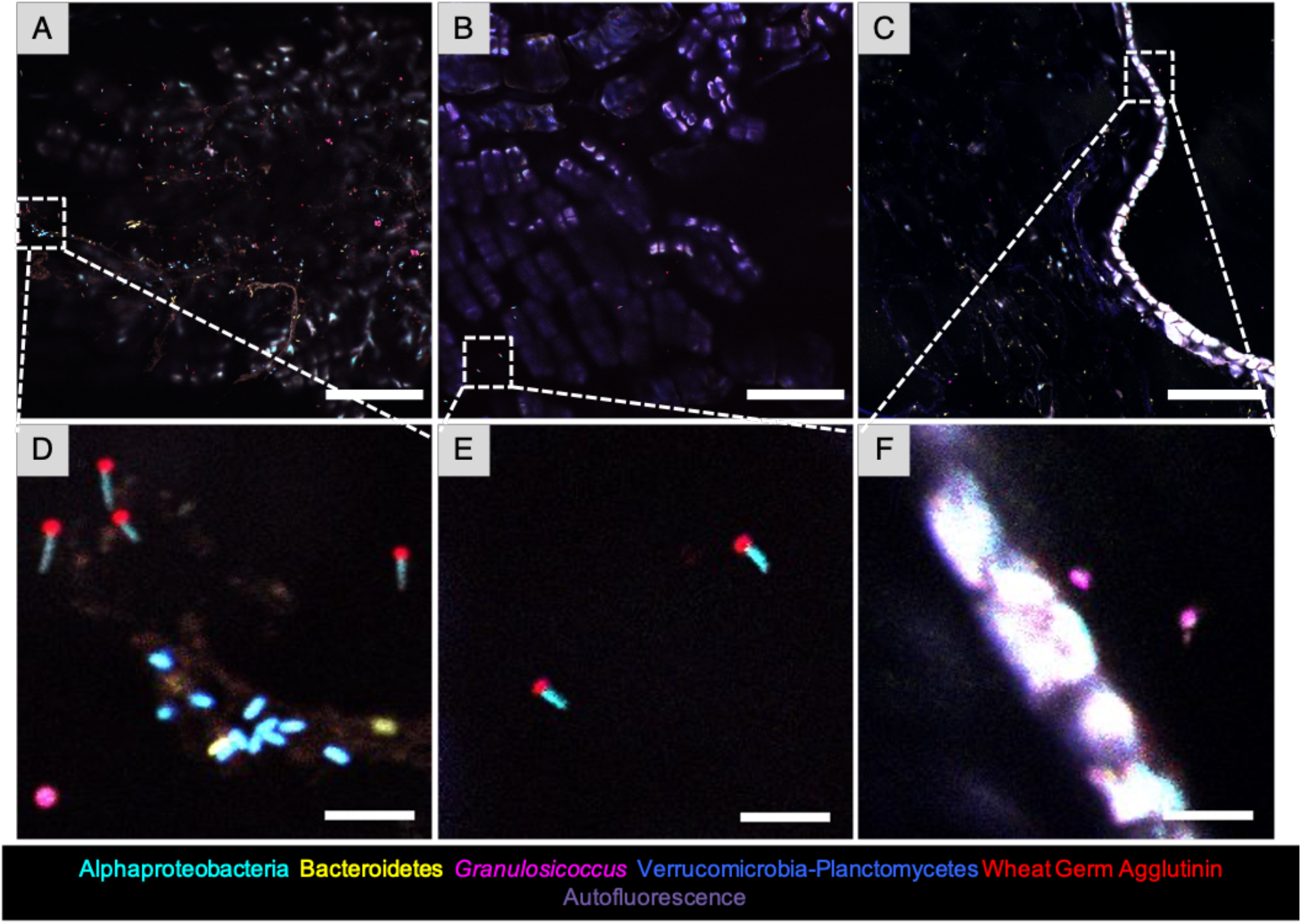
Low microbial density on declining population of kelp in Squaxin Island. (A) and (B) whole mount FISH showing sparse bacteria on kelp surface. (C) Cross-section in which no dense biofilm is observed on the surface, but a few bacteria were visible. Strong autofluorescence of kelp cells is observed. (D), (E) and (F) are enlarged images of the dashed squares in (A), (B), and (C), respectively. Scale bar = 20 μm (A-C); 5 μm (D-F).

### Bacterial density differences between samples from healthy compared to declining kelp

In addition to the samples collected on the outer Pacific Ocean site of Tatoosh Island, we imaged the microbial biofilm of kelp collected at Squaxin Island, located in Southern Puget Sound. Although the same major taxa are present on *N. luetkeana* from both collection sites, their relative abundances differed: Alphaproteobacteria accounted for the bulk (∼75%) of the microbial community in 16S rRNA gene sequences from Squaxin samples (Fig. 1). Imaging revealed dramatically lower density of microbes in Squaxin samples (Fig. 8) compared to the tip of Tatoosh blades and similar densities to the Tatoosh base samples. Interestingly, kelp blades at Squaxin Island were sampled during the same week as the late June Tatoosh kelp (Table S1, June 21 vs. June 25), yet the Tatoosh kelp (Fig. 4A) have a much greater density of bacterial cells than the Squaxin Island kelp. While it is difficult to infer an exact comparison, given that the Squaxin kelp blades were sampled from the middle of the blade while the Tatoosh kelp communities were sampled at the base and tip of the blade, the estimated age of sampled tissue from Squaxin kelp was approximately 2 months, similar to that of the tissue collected from the tip of the kelp blade on Tatoosh Island (Table S1). Thus, samples collected at the base of the Tatoosh blades that were less than 1 week old had similar microbial densities to these 2-month-old biofilms on kelp from Squaxin Island.

### Micron-scale spatial arrangement

Using image analysis with *daime* (Daims et al. 2006), we counted a mean of 4025 *Granulosicoccus* cells and 2445 Verrucomicrobia-Planctomycetes cells per image on 8 *Nereocystis* individuals (Table 2). Overall densities of *Granulosicoccus* averaged 89,056 cells per mm^2^ and as high as 269,049; the Verrucomicrobia-Planctomycetes probe identified an average of 54,100 cells per mm^2^, with a maximum of 94,530. These numbers are likely underestimates of the total colonization of the kelp surface, as they represent only the cells visible in a single focal plane. Cell size was comparable with the Verrucomicrobia-Planctomycetes measuring 0.42 and *Granulosicoccus* measuring 0.47 μm in diameter across all 8 individuals (Table 2).

**Table 2.** Cell size, abundance and spatial correlation using linear dipole analysis in *daime* across 8 individuals (n=2-13 replicates each) in images 212.55 μm on a side. Means (SD) are given. Cell abundances were positively correlated within and among the two species, indicating clumped distributions. The distribution of correlations with distance are shown in FigS3.

| Probe | Cell size in $\mu\text{m}$ | Cell Abundance | Distances of spatial autocorrelation | Distances of spatial cross correlation |
| --- | --- | --- | --- | --- |
| *Granulosicoccus* | 0.47 (0.23) | 4025.3 (3416.7) | <9 $\mu\text{m}$ | - |
| Verrucomicrobia | 0.42 (0.16) | 2445.3 (1142.2) | <7 $\mu\text{m}$ | - |
| *Granulosicoccus* vs. Verrucomicrobia | - | - | - | <5 $\mu\text{m}$ |

Linear dipole analysis can be used to calculate the spatial correlation between two taxa, or between a taxon and itself, over a range of distances. We carried out linear dipole analysis in *daime* to analyze within-taxon and between taxon associations for the genus *Granulosicoccus* and the Verrucomicrobia-Planctomycetes (hereafter referred to as simply Verrucomicrobia) for each of 8 kelp individuals. Results indicated that both *Granulosicoccus* and Verrucomicrobia cells were positively autocorrelated spatially at distances less than 10 um (Table 2, Fig. S3). When we quantified spatial covariance for both taxa, their co-occurrence was observed at distances less than 5 μm and peaked at 1.61 μm (Table 2, Fig. S3).

While these correlations may reflect a tendency of these specific cell types to localize close to one another, they could also result from overall patchiness in colonization of the frond, or from imaging or preparation artifacts (e.g. from compression of the samples between slide and coverslip, or conversely from the sample being not entirely flat in the plane of focus).

## Discussion

### Comparison of the *Nereocystis* microbiota to other microbiomes

CLASI-FISH reveals the microbiota of *Nereocystis* to be dense and complex. The bacterial community growing on or within macroalgal fronds has been visualized by other investigators using FISH with one or a few taxa (Tujula et al. 2006, 2010; Bengtsson and Ovreas 2010), showing a biofilm composed of cocci and filaments and highlighting the distribution of individual taxa. The benefit of CLASI-FISH is the ability to identify the major groups simultaneously and assess their relative distribution and potential for direct spatial interaction. The probe set we employed provides rather coarse resolution, with most of the targeted taxa visualized at the phylum or class level; nonetheless we visualized micron-scale interaction among these diverse taxa and our results lay out a framework that can be furthered by future experiments using more complex and specific probe sets.

The microbiota visualized here is notable for its high density but moderate thickness. In the most densely colonized samples, our images show a confluent biofilm several cells thick. With a typical microbial diameter of 0.5 μm, this confluent biofilm corresponds to a density of 10^8^-10^9^ cells/cm^2^, comparable to the density of 10^7^-10^8^ cells/cm^2^ measured for the brown alga *Fucus vesiculosus* (Stratil et al. 2013). Other marine organisms have a lower surface colonization density; for example, the density of microbes in coral mucus is estimated at only 10^5^-10^6^ per cm^3^ (Garren and Azam 2010; Glasl et al. 2016), on the same order of magnitude as the density in surrounding seawater. Compared to the communities that grow on the human tongue and teeth, the kelp microbiota has similar complexity, but its thickness, in the range of 10 microns, is limited compared to the potentially hundreds of microns in thickness achieved by the microbiota of the human mouth (Zijnge et al. 2010, Mark Welch et al. 2016, Wilbert et al. 2019 in press).

### Functional implications of the kelp microbiota

The functional importance of the dense microbial layer that we have revealed through CLASI-FISH is relatively unknown, but the position of the microbial layer at the interface between the host tissue and the surrounding water column suggests the possibility for important consequences for biofouling, access of the host organism to light and nutrients, and metabolic exchange. Mucus production by kelps may play a critical role in providing structure for surface-associated microbes. Potential benefits of these microbes to the host may include generation of antibacterial compounds that protect the host against fouling and pathogens (Rao et al. 2007; Egan et al. 2013, Michelou et al. 2013, Tebben et al. 2014, Lee et al. 2016) or competitors (Barott and Rohwer 2012). In turn, microbes likely benefit from a predictable source of dissolved organic matter and a persistent substrate for colonization. Possible functional interactions between macroalgae and their epibionts, both detrimental and potentially beneficial, have been the subject of several recent reviews (Wahl et al. 2012; Ramanan et al. 2016; Singh and Reddy 2016; Florez et al. 2017).

The limited thickness of the microbial biofilm that we imaged raises the question of whether this biofilm is thin enough so as to permit unattenuated light penetration to the kelp itself and raises the question of what mechanisms may exist by which the thickness of the biofilm may be limited. Is there a dynamic process of biofilm loss via host shedding of the mucous coating to which the biofilm may adhere, followed by re-growth of the biofilm? Alternatively, is biofilm thickness limited by the intrinsic rate of growth of the microbes or their accretion from the water column, or by grazing of the biofilm by micro- or macro-invertebrates? Interestingly, the *Granulosicoccus sp*. genome contains 30 flagella genes (Kang et al. 2018), leading to the possibility that this bacterium is motile, which would provide a mechanism for its early colonization of the kelp tissue from the seawater as well as its high abundances on the kelp surface.

Imaging reveals close association, at micrometer scales, of different microbial taxa with one another and with the host, a spatial organization that creates the conditions necessary for metabolic exchange among microbes (Seth and Taga 2014) and between host and microbiota. While recent studies have described microbial communities in association with kelp through genomics (Michelou et al. 2013, Lemay et al. 2018, Qiu et al. 2019), the metabolic role of the microbes relative to the host has yet to be clarified. Nutrient exchanges between host and microbes are functionally significant in phytoplankton (e.g. Amin et al. 2015). *Nereocystis* at Tatoosh Island release ∼20% of fixed carbon into the surrounding seawater as dissolved organic carbon (DOC) (Weigel and Pfister, in review), a quantity consistent with DOC release estimates for other kelp species (Abdullah & Frederiksen 2004, Reed et al. 2015). By living on the kelp surface, biofilm microbes are presented with a consistent and labile metabolic resource, in addition to the structural kelp polymers that kelp microbes can degrade (Bengtsson et al. 2011).

The release of carbohydrate exudates likely favors heterotrophic microbial metabolisms, and the *Granulosicoccus sp*. sequence variant in this study shares 97% sequence identity to *Granulosicoccus antarcticus* (Lee at al. 2007), which is a heterotrophic microbe that contains urease and both nitrate and nitrite reductase genes (Kang et al. 2018). As a heterotroph, *Granulosicoccus sp*. likely takes advantage of the abundant DOC, while nitrogen transformation genes suggest a potential for nitrogen metabolisms that may impact the host kelp. Likewise, studies of microbial nutrient transformation in near-shore waters of Tatoosh Island showed that these microbial nitrogen metabolisms were strongest in association with the surfaces of a red alga, *Prionitis sternbergii*, rather than in seawater or associated with inert substrates (Pfister and Altabet 2019). This finding suggests that epibiont communities on algae are enriched for microbes carrying out ammonium oxidation and nitrate reduction, both of which might serve to retain and recycle dissolved inorganic nitrogen (DIN) near the surface of the alga.

### Low density of microbiota and high fraction of Alphaproteobacteria on declining population of kelp

Shifts in microbial composition between healthy and stressed macroalgae have been reported (Marzinelli et al. 2015), but the low density of bacteria on the kelp from Squaxin Island was unexpected, as we had initially assumed that a population in decline would be more likely to be overrun with microbes than nearly devoid of them. The majority of the bacterial epibionts at Squaxin were shown by both sequencing and imaging to be Alphaproteobacteria, most likely represented by the single highly-abundant ASV in the sequencing data identified as a member of the Hyphomonadaceae. Alphaproteobacteria from the family Rhizobiales produce unipolar adhesins which are essential for cell-cell adhesion, biofilm formation and effective root colonization (Fritts et al. 2017; Williams et al. 2008). The exopolysaccharide N-acetylglucosamine, synthesized by bacterial cells, plays an important role in biofilm formation in *Staphylococcus aureus* (Izano et al. 2008, Lin et al. 2015) and *Escherichia coli* (Wang et al. 2004). Interestingly, the consistent presence of N-acetylglucosamine or N-acetylmuramic acid residues at one end of *Alphaproteobacteria* cells suggests that it may be involved in cell adhesion to the kelp mucous layer. The high relative abundance of Alphaproteobacteria in Squaxin samples, in which the observed bacteria density is low, might be related to a similar attachment strategy that would allow them to attach to the kelp surface more permanently or more readily than other members of the microbiota.

## Materials and Methods

### Sample collection and 16S rRNA gene sequencing

Photosynthetic blade tissues of *Nereocystis luetkeana* were sampled at five time points spaced roughly 2 weeks apart (11 June – 22 August 2017) on Tatoosh Island, Washington in the United States (48°23’37.0”N 124°44’06.5”W). At each time point, two tissue samples (2 × 1 cm^2^) were collected from a single blade – one at the basal meristem, roughly 2 cm from where the blade connects to the stipe, to capture recently produced tissue (∼24 to 48 hours old) and another near the apical end of the blade tip to sample older tissue (weeks to months old). Samples were collected from different kelp individuals at each date, and *n* = 2 or 3 samples from each date were selected for imaging. Total blade length and linear blade growth rates were measured to approximate the age of tissues sampled (Weigel and Pfister 2019). Kelp blade tissues samples were also collected from Squaxin Island, in the Southern Puget Sound (47°10’38.7”N 122°54’42.2”W) on 21 June 2017. At this site, kelp tissue samples were collected from the middle of the kelp blade, and *n* = 3 samples were selected for imaging. Kelp blade tissues for CLASI-FISH and 16S rRNA sequencing were collected together from adjacent locations on the kelp blade. Samples collected for CLASI-FISH were preserved in 95% ethanol and stored at −20°C, while samples for 16S rRNA had no preservatives and were temporarily frozen at −20°C until they were shipped to −80°C. Details of DNA extraction, 16S rRNA gene sequencing, and sequence analysis are contained in Weigel and Pfister (2019).

### Probe design for *Granulosicoccus* and Probe set validation

Probes for genus *Granulosicoccus* were designed based on sequencing results. An alignment of all *Granulosicoccus* V4 fragment sequences and most abundant ASVs from all the other taxa was performed using Geneious 11.1.3. Alignment was reviewed manually to look for candidate regions for probe design. We selected sequence regions which match all the *Granulosicoccus* sequences and did not match any of the other taxa. Probe hybridization efficiency was evaluated using mathFISH (Yilmaz et al., 2011). Two probes were designed:Gran670 (5’-CACCGCTACACCCGGAATTCCGC-3’) and Gran737 (5’-TCAGCGTCAGTATTGTTCCAGA-3’).

To validate the probes for specificity, we applied the set of 7 probes simultaneously to pure cultures and hybridized and imaged under the same conditions as kelp samples (Fig. S1). Probes used in this study are listed in Table 1.

### Bacterial strains and growth conditions

*Granulosicoccus coccoides* DSM 25245 and *G. antarcticus* DSM 24912 were cultured in Bacto Marine Broth media (DIFCO 2216) (pH 7.5 and 7 for *G. coccoides* and *G. antarcticus*, respectively). Cultures were incubated with agitation (180 rpm) at 25^°^C. Two- and 7-days cultures were fixed with 2% PFA on ice for 90 min, washed and transferred to 50% ethanol. Fixed cells were stored at −20^°^C until use.

### Embedding and sectioning for imaging

For methacrylate embedding, kelp samples stored in 95% ethanol at −20°C were placed in 100% ethanol for 30 min followed by acetone for 1 hour, infiltrated with Technovit 8100 glycol methacrylate (EMSdiasium.com) infiltration solution 3 h replacing for fresh solution every hour and followed by a final infiltration overnight at 4^°^C. Samples were then transferred to Technovit 8100 embedding solution and solidified for 12 h at 4 °C. Blocks were sectioned to 5 μm thickness using a Leica microtome (RM2145) and applied to Ultrastick slides (Thermo Scientific). Sections were stored at room temperature until CLASI-FISH was performed.

### Combinatorial labeling and spectral imaging fluorescence *in situ* hybridization (CLASI-FISH)

CLASI-FISH microscopy was carried out on a subset of samples analyzed previously by 16S rRNA sequencing and reported in Weigel and Pfister (2019).

We used two methods to visualize the spatial structure of the kelp microbiota: whole-mount-agarose preparations and methacrylate sections. When possible, we used pieces of the same kelp sample for both methods. For the whole-mount-agarose method one piece of kelp was placed on a slide and 50 μl of 1% low-melting-point agarose was dropped on it and the sample-allowed to cool on ice for 10 min before the CLASI-FISH procedure. Methacrylate sections were subject to CLASI-FISH directly on slides.

Hybridization solution [900 mM NaCl, 20 mM Tris, pH 7.5, 0.01% SDS, 20% (vol/vol) formamide, each probe at a final concentration of 2 μM] was applied to kelp pieces and incubated at 46 °C for 2 h in a chamber humidified with 20% (vol/vol) formamide. Whole-mount-agarose preparations were maintained in horizontal position and were washed with 100 μl of pre-warmed wash buffer (215 mM NaCl, 20 mM Tris, pH 7.5, 5mM EDTA) five times at RT followed by three washes with 500 μl of wash buffer at 48°C for 5 min each. The hybridization conditions were the same for methacrylate sections, but washing was carried out by incubating the slides in 50 ml of washing buffer for 15 minutes at 48^°^C. Samples were then incubated with wheat germ agglutinin (20 μg ml^−1^) conjugated with Alexa Fluor 680 at room temperature for 30 min in the dark. Agarose-coated samples were washed with 100 μl of sterile cold water three times. Excess agarose was cut off with a disinfected razor. Methacrylate sections were washed by dipping the slide into 50 ml of ice-cold water to remove excess salt. Samples were mounted in ProLong Gold antifade reagent (Invitrogen) with a #1.5 coverslip and cured overnight in the dark at room temperature before imaging.

### Image acquisition and linear unmixing

Spectral images were acquired using a Carl Zeiss LSM 780 confocal microscope with a Plan-Apochromat 40X, 1.4 N.A. oil immersion objective. Images were captured using simultaneous excitation with 405, 488, 561, and 633 nm laser lines. Linear unmixing was performed using ZEN Black software (Carl Zeiss) using reference spectra acquired from cultured cells hybridized with Eub338-I probe labeled with one of the 6 fluorophores in the probe set and imaged as above. Unmixed images were assembled and false-colored using FIJI software (Schindelin *et al.*, 2012).

### Spatial Analysis of CLASI-FISH images

We used the software *daime* (Daims et al. 2006) to analyze the spatial structure and the size of the cells identified with CLASI-FISH on 8 separate *Nereocystis* individuals (n = 2 to 13 images each) collected between 11 Jun to 22 Aug 2017. We imported images that were 212.55 μm on a side for each probe separately as TIFF files, enhanced the contrast manually, and used automatic 2D segmentation with the RATS-L thresholding algorithm to select and count cells. Very small objects were interpreted to be noise and were manually removed using the object editor. We quantified spatial distribution using ‘scan whole reference space’ to a distance of 50 μm and recorded the correlation at every 0.15 μm using the 2D linear dipole algorithm.

## Acknowledgements

Funding was provided by NSF award 1650141 (to JMW) and the NOAA-COCA program through NA16OAR431055 (to CAP). BLW was supported by a National Geographic Society Early Career Grant, a Phycological Society of America Grants-in-Aid of Research, a Department of Education GAANN fellowship, and a travel award from the Committee on Evolutionary Biology at the University of Chicago. LJ was supported in part by NSF REU award 1659604, “Biological Discovery in Woods Hole at the Marine Biological Laboratory.” We thank the Makah Tribal Nation for access to Tatoosh Island, and H. Berry, T. Mumford and the Washington DNR for boat logistics at Squaxin Island. T. Bowyer, K. Miranda, A.M. Wootton, J.B. Wootton, and J.T. Wootton all helped in the field. We thank Tracy Mincer, Linda Amaral-Zettler and Erik Zettler for providing microbial cultures for probe testing.

The funders had no role in study design, data collection and interpretation, or the decision to submit the work for publication.

