## Supplemental figures and table for "Spatial organization of the kelp microbiome at micron scales"

### Supplemental Material

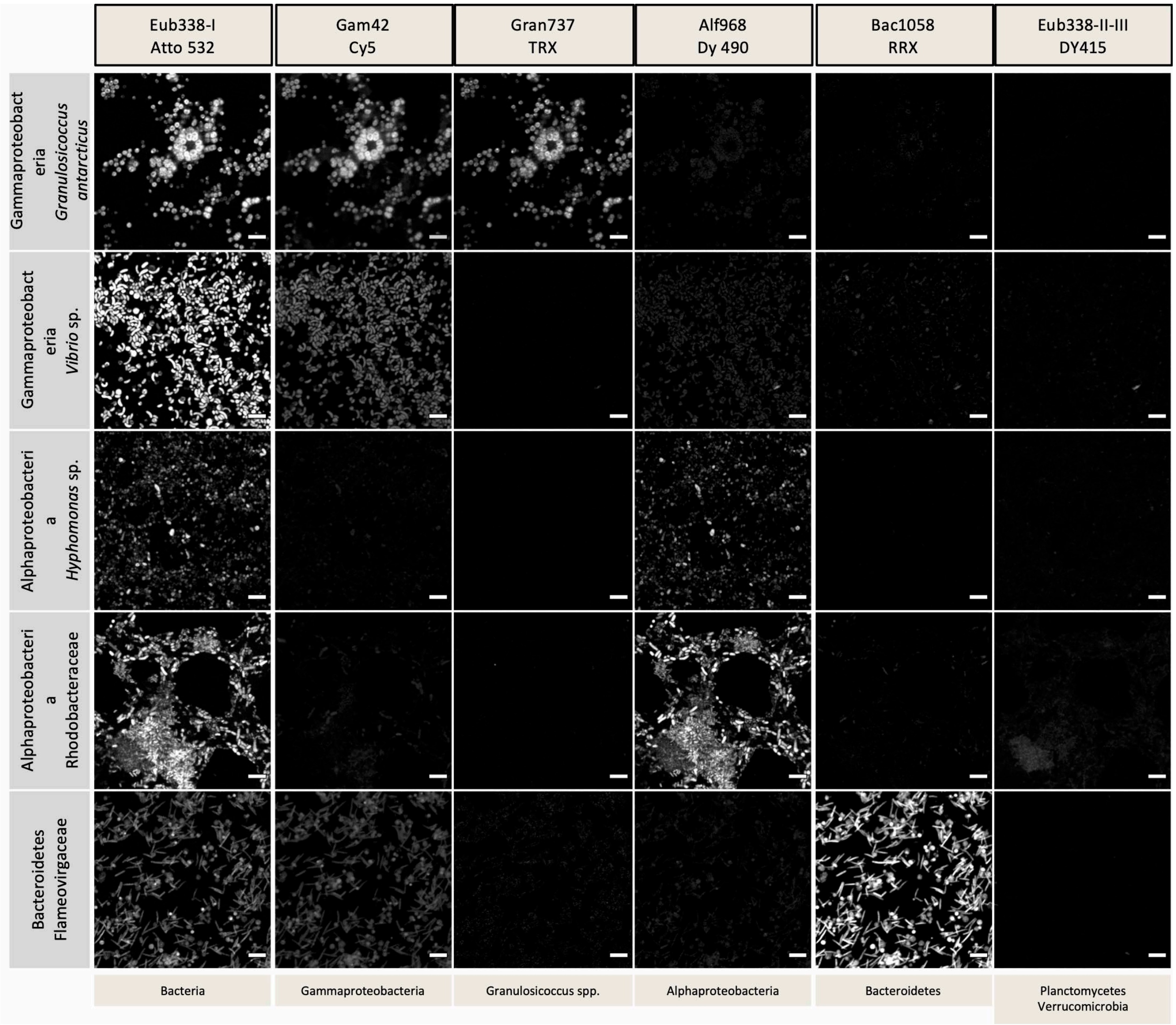


**Figure S1.** Validation matrix of a set of 7 probes, each labeled with a distinct fluorophore except for Eub338-II and Eub338-III which were both labeled with the fluorophore Dy415. To validate the probes for specificity, we applied the set of probes to pure cultures, hybridized and imaged under the same conditions as kelp samples. Results show that each specific probe hybridized with its expected target taxa; some cross-reactions are visible (e.g., Gam42a probe with Bacteroidetes cells) but are faint relative to hybridization of those same cells with the probe targeting them (e.g., Bac1058 probe with Bacteroidetes cells). Probe name is shown at top of each column. Bacterial culture names are shown in left column. Target taxon for each probe is shown in row in the bottom. Scale bars= 5 µm.

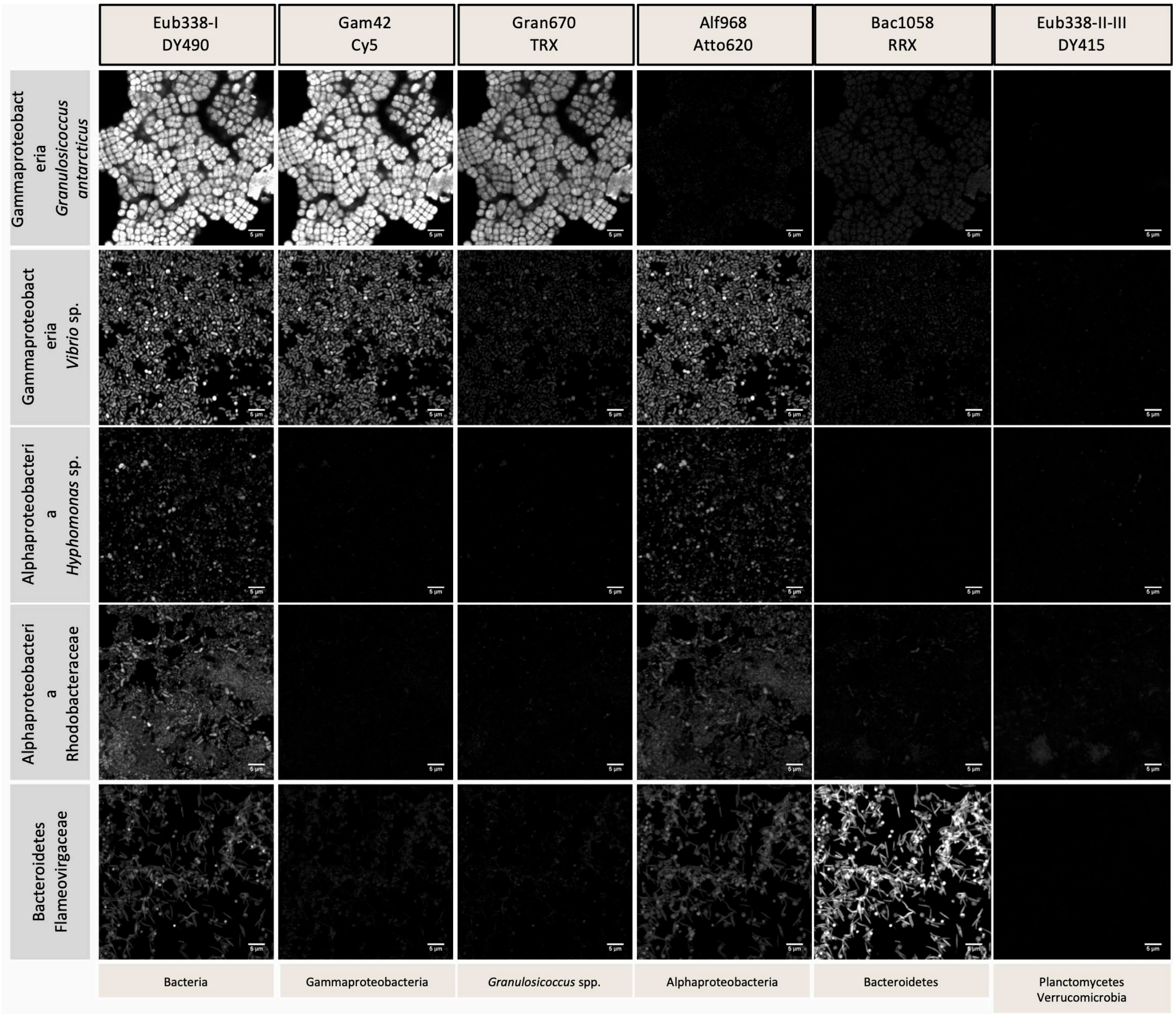


**Figure S2.** Validation matrix of a set of 7 probes, each labeled with a distinct fluorophore except for Eub338-II and Eub338-III which were both labeled with the fluorophore Dy415. To validate the probes for specificity, we applied the set of probes to pure cultures, hybridized and imaged under the same conditions as kelp samples. Results show that each specific probe hybridized with its expected target taxa; some cross-reactions are visible (e.g., Gam42a probe with Bacteroidetes cells) but are faint relative to hybridization of those same cells with the probe targeting them (e.g., Bac1058 probe with Bacteroidetes cells). Probe name is shown at top of each column. Bacterial culture names are shown in left column. Target taxon for each probe is shown in row in the bottom. Scale bars= 5 µm.

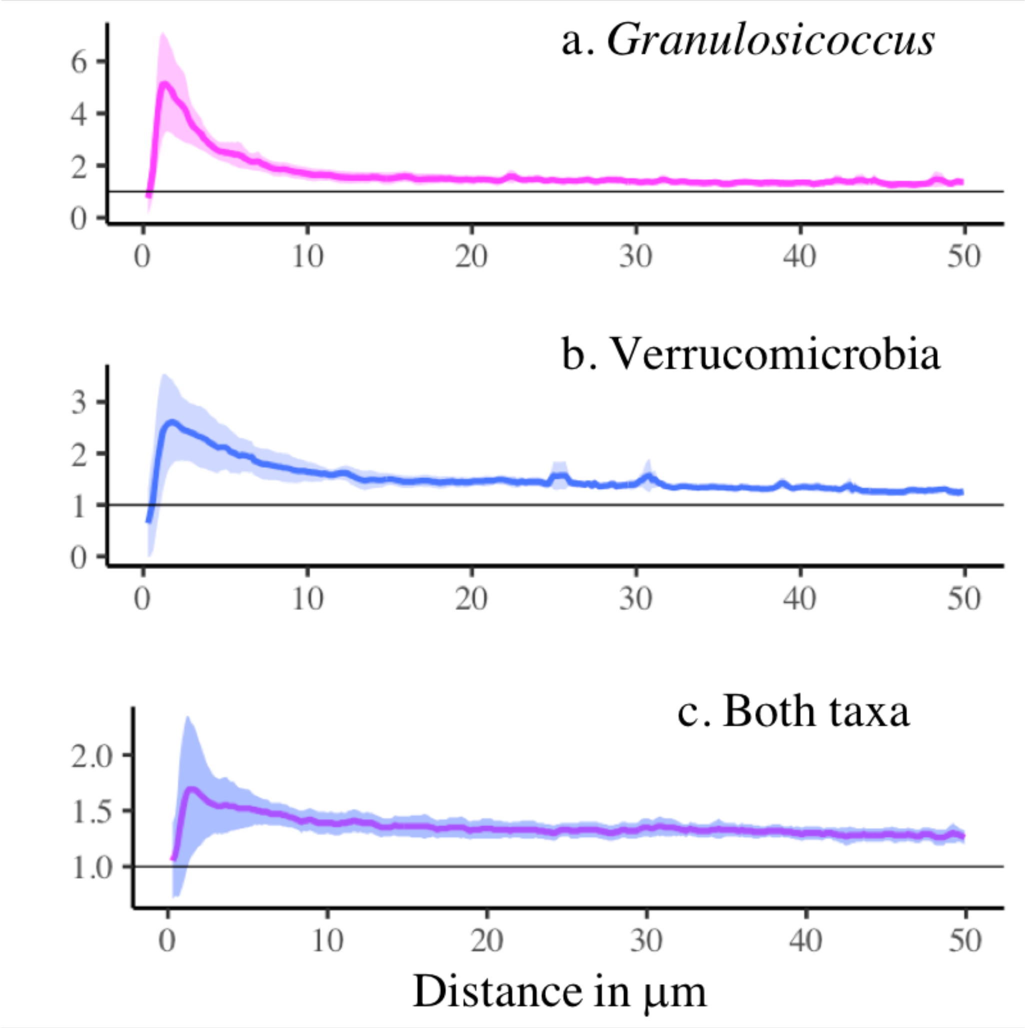


**Figure S3.** The autocorrelation in a. Granulosicoccus and b. Verrucomicrobia based on spatial analysis with *daime* showing strong association in each taxon. *Granulosicoccus* cells were correlated to each other at 0.59 µm and beyond with a maximum at 1.32 µm. Verrucomicrobia cells were correlated to each other at 1.17 µm and beyond with a maximum at 1.19 µm. The spatial correlation between the 2 taxa was also significant starting at 1.32 µm and peaking at 1.61 µm. The mean (solid line) and 95% confidence interval (ribbon) are shown based on 8 individual kelp with 2 to 13 censuses on each.

**Table S1**. Samples of *N. luetkeana* kelp used for CLASI-FISH.

| Sample ID | Blade sample type | Collection date | Collection location | Total blade length (cm) | Estimated age tissue (days) | 16S Sequence Count | % Bacterial Sequences |
|---|---|---|---|---|---|---|---|
| 63-B | Base | 6/11/17 | Tatoosh | 124 | < 7 | 27,927 | 1.5 |
| 63-A | Tip | 6/11/17 | Tatoosh | 124 | 62 | 42,113 | 81.8 |
| 61-B | Tip | 6/11/17 | Tatoosh | 95 | 48 | 51,654 | 89.6 |
| 81-B | Base | 6/25/17 | Tatoosh | 87 | < 7 | 32,046 | 3.7 |
| 81-A | Tip | 6/25/17 | Tatoosh | 87 | 44 | 51,006 | 89.7 |
| 82-A | Tip | 6/25/17 | Tatoosh | 131 | 66 | 50,981 | 89.6 |
| 128-A | Tip | 7/10/17 | Tatoosh | 102 | 51 | 48,165 | 88.1 |
| 132-A | Tip | 7/10/17 | Tatoosh | 156 | 78 | 73,121 | 89.6 |
| 158-B | Base | 7/24/17 | Tatoosh | 146 | < 7 | 31,388 | 2.3 |
| 158-A | Tip | 7/24/17 | Tatoosh | 146 | 73 | 61,338 | 97.0 |
| 271-B | Base | 8/22/17 | Tatoosh | 72 | < 7 | 14,014 | 1.5 |
| 271-A | Tip | 8/22/17 | Tatoosh | 72 | 36 | 49,759 | 90.3 |
| 71 | Middle | 6/21/17 | Squaxin | 268 | 67 | 41,277 | 34.3 |
| 72 | Middle | 6/21/17 | Squaxin | 266 | 67 | 39,558 | 7.6 |
| 73 | Middle | 6/21/17 | Squaxin | 302 | 76 | 42,700 | 10.1 |
